# 2-D Neural Geometry Underpins Hierarchical Organization of Sequence in Human Working Memory

**DOI:** 10.1101/2024.02.20.581307

**Authors:** Ying Fan, Muzhi Wang, Nai Ding, Huan Luo

**Author notes:** Correspondence (N.D.) (H.L.).

## Abstract

Working memory (WM) is constructive in nature. Instead of passively retaining information, WM reorganizes complex sequences into hierarchically embedded chunks to overcome capacity limits and facilitate flexible behavior. To investigate the neural mechanisms underlying hierarchical reorganization in WM, we performed two electroencephalography (EEG) and one magnetoencephalography (MEG) experiments, wherein humans retain in WM a temporal sequence of items, i.e., syllables, which are organized into chunks, i.e., multisyllabic words. We demonstrate that the 1-D sequence is represented by 2-D neural representational geometry in WM arising from parietal-frontal regions, with separate dimensions encoding item position within a chunk and chunk position in the sequence. Critically, this 2-D geometry is observed consistently in different experimental settings, even during tasks discouraging hierarchical reorganization in WM and correlates with WM behavior. Overall, these findings strongly support that complex sequences are reorganized into factorized multi-dimensional neural representational geometry in WM, which also speaks to general structure-based organizational principles given WM’s involvement in many cognitive functions.

## Introduction

Working memory (WM) is constructive in nature^1–3^. Instead of passively storing information, the WM system actively builds new representations to fulfill task-specific goals and deal with various contexts^4–8^. In fact, most high-level cognitive functions, including language, motor control, reasoning, and planning, involve reorganizing preexisting elements in ways that differ from their original formats^9–14^. For example, when memorizing a sequence of items such as phone numbers or words, we group them into chunks^15^. When forming episodic memory of a movie or a novel, a continuous stream of events tends to be reorganized into hierarchically embedded episodes^16^. Reconfiguration of successive items into hierarchical embedded chunks or schemas is a major memory operation to overcome capacity limits, promote flexible goal-directed processing, and facilitate generalization in novel scenarios^17–20^.

Previous behavioral evidence also supports the hierarchical reorganization of sequences in WM, by revealing transposition WM errors for items occupying the same position across different chunks^21–25^. Based on these findings, computational models further propose that each item in a hierarchical sequence would be represented based on two indexes: a global index indicating its affiliated chunk order and a local index denoting its position within a chunk^26–28^. However, the neural implementation of this hierarchical reorganization in WM remains unknown. Answers to this question would shed light on the neural mechanism of WM operation and provide substantial insights into a wide range of fields given WM’s involvement in almost any cognitive function.

To achieve abstract hierarchical organization, structure and content are proposed to be represented in a disentangled manner, known as factorization^29,30^. This view has been proposed in computational models for sequence memory^31,32^ and supported by recent empirical findings^33–36^. For instance, prefrontal cortex (PFC) recordings in monkeys showed that when retaining a sequence of spatial locations in WM, each ordinal rank in the sequence occupies a subspace in multidimensional neural space, regardless of its content, i.e., the memorized location^37^. Furthermore, memorized content and sequence structure are reactivated by different triggering events during auditory WM retention, indicating their dissociated neural formats^38,39^. In light of these findings, we postulate that WM contains factorized representations of hierarchical structure and content items, and here we aimed at examining the neural representation of abstract hierarchical structure regardless of the items being attached. What types of neural representation could support the abstract hierarchical structure underlying sequence organization in WM? One straightforward possibility is a 1-D chunked format, in which items within a chunk have compressed representational distance to each other, compared with items in different chunks (Figure 1B, right middle). In contrast, however, here we propose that the hierarchical structure could also be implemented by 2-D neural geometry whereby global and local ranks are separately encoded as separate dimensions spanning a 2-D space (Figure 1B, right lower). In other words, a 1-D sequence embedded in hierarchical structures is neurally represented along two dimensions. Our hypotheses are motivated by previous findings: First, behavioral and modeling work propose that each item in a hierarchical sequence is reorganized via two indexes: a global index and a local index^21–23,25–28^. Second, monkey recordings show that ranks of simple non-hierarchical sequences are encoded in near-orthogonal neural manifolds in WM ^37^. This geometry operation strategy, also observed in other fields, is suggested to minimize interference between cognitive variables^40–42^. Moreover, different from most studies examining the encoding period when items are presented^43,44^, here we specifically focused on the maintaining period which would better reflect the internal organizational principle of the WM system.

**Figure 1.**
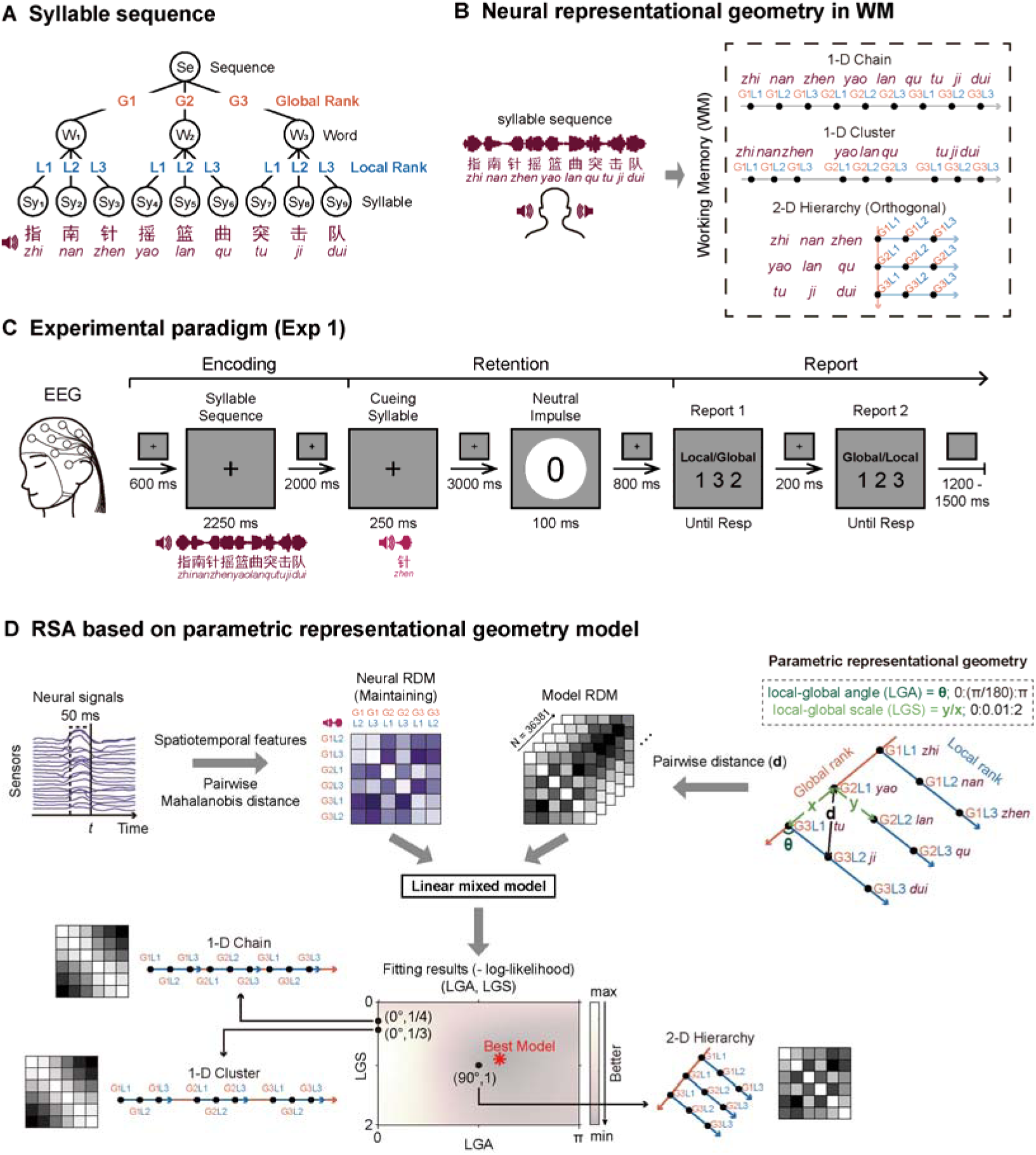
Representational geometry models of syllable sequences in WM. (A) Example of hierarchically organized Chinese syllable sequence. The 9-syllabel sequence comprises three trisyllabic words, with each syllable associated with one global rank (orange; G1, G2, G3) and one local rank (blue; L1, L2, L3). The sequence was presented iso-rhythmically at 4 Hz and randomly generated per trial. (B) Three hypotheses (1-D Chain, 1-D Cluster, 2-D Hierarchy) characterizing neural representational geometry of the syllable sequence in WM. 1-D Chain: syllables in the sequence reside evenly along a 1-D dimension. 1-D Cluster: same as 1-D Chain but with shorter within-word neural distance. 2-D Hierarchy: two separate axes to encode global and local ranks of a sequence, with its strongest form as the 2-D Orthogonal hypothesis positing orthogonalization of local and global dimensions. (C) Experiment 1 paradigm. Subjects retained a 9-syllabe sequence in WM with EEG recordings. During retention, one of the 9 syllables was presented (cuing syllable), followed by a neutral visual impulse. Subjects serially reported the global and local rank of the cueing syllable during report (order balanced across trials). (D) Illustration of representational dissimilarity analysis (RSA) based on parametric representational geometry model. Neural RDM (left, purple): multivariate Mahalanobis neural distance between rank-specific responses regardless of the attached syllable were calculated using a sliding 50-ms time window. Model RDM (right, grey): a parametric representational geometry model with two free parameters: local-global angle (LGA) and local-global scale (LGS). LGA: angle between local and global dimensions. LGS: ratio of unit length between local and global dimensions. A series of hypothetical neural representational geometries and corresponding model RDMs were constructed by sampling combination of the two parameters (LGA: 0°:1°:180°; LGS: 0:0.01:2). Lower panels: each model RDM was regressed to neural RDM (linear mixed model), resulting in model fitting performance (negative log-likelihood) as a function of LGA and LGS. The one with the lowest negative log-likelihood value was regarded as the best model (red asterisk). Note all the three neural representational geometry hypotheses could be captured using the parametric representational geometry model with different LGA and LGS values (1-D Chain: LGA = 0°, LGS = 0.33;1-D Cluster: LGA = 0°, LGS = 0.25; 2-D Orthogonal: LGA = 90°, LGS = 1).

To test the hypothesis, we performed two electroencephalography (EEG) and one magnetoencephalography (MEG) experiments that asked subjects to retain a series of syllable sequences, which were hierarchically organized into words and multi-word sequences, and perform a rank recalling task. We examined the neural geometry of the syllable sequence in WM using an innovative parametrical representational similarity analysis (RSA) approach. We demonstrate a 2-D factorized representation of the syllable sequence, with separate dimensions for the local rank (position of a syllable within a word) and the global rank (position of a word within a sequence). Critically, this 2-D neural geometry, originating from parietal-frontal brain region, is observed consistently in different stimulus settings and tasks, even when the task discourages hierarchical structures, and correlates with memory behavior. Overall, these results support that the WM system can reorganize a complex linear sequence into a 2-D factorized neural representation to reveal the underlying hierarchical structure.

## Results

### Representational geometry models of syllable sequences in WM

Hierarchically organized syllable sequences were used to assess the neural representational geometry of sequence retained in WM (Figure 1A). In Experiment 1, each trial presented 9 Chinese syllables at a constant rate of 4 Hz, which grouped into three trisyllabic words (different words in different trials). Accordingly, each syllable is associated with two ordinal ranks – a global rank (index of word in the sequence) and a local rank (index of syllable within a word). Human subjects were instructed to retain the sequence in WM. One of the 9 syllables was presented during retention as the cueing syllable and subjects serially reported the global and local ranks of the cueing syllable during retrieval (balanced order across trials) (Figure 1C).

Here, we focus on the neural representational geometry of sequence in WM during the retention period, and formulate three hypotheses (Figure 1B). The 1-D Chain hypothesis postulates even spacing of the neural representation of each syllable along an axis, mirroring their presentation order. The 1-D Cluster hypothesis extends the 1-D Chain hypothesis by additionally considering the grouping of syllables into words and postulates that syllables within a word have shorter neural representational distances than those spanning word boundaries. In contrast, the 2-D Hierarchy hypothesis postulates separate neural axes for local and global ranks. The strongest form of the 2-D Hierarchy hypothesis is that the two dimensions are orthogonal to each other. Under this condition, referred to as the 2-D Orthogonal hypothesis, there is no interference between global-rank and local-rank representations.

We quantified the three hypotheses using a parametric representational geometry model with two free parameters, i.e., the local-global angle (LGA) and the local-global scale (LGS) (Figure 1D, right). The LGA denotes the angle between the local and global dimensions. The 1-D Chain and Cluster hypotheses predict a 0° LGA, i.e., local and global ranks occupy the same representational dimension. The 2-D Hierarchy hypothesis predicts a non-zero LGA and the 2-D Orthogonal hypothesis predicts a 90° LGA. The other parameter, LGS, denotes the ratio of representational distance between consecutive units along the local and global dimensions. The LGS is 1/3 according to the Chain hypothesis and should be lower than 1/3 according to the Cluster hypothesis, which assumes compressed within-word distance. The 2-D Hierarchy or Orthogonal hypothesis does not constrain the LGS. However, a LGS lower than 1 indicates that global rank is better discriminated by the neural response than the local rank, and vice versa. Therefore, when the neural data was fitted using the representational geometry model, the fitted parameters could reveal whether the neural representation of the syllable sequence was consistent with 1-D Chain (LGA = 0°, LGS = 1/3), 1-D Cluster (LGA = 0°, LGS < 1/3), or 2-D Hierarchical hypothesis (LGA ≠ 0° and LGA = 90° for the 2-D Orthogonal hypothesis).

We employed a representational dissimilarity analysis (RSA) to investigate which parameters of the representational geometry model best fit the neural data during WM retention. We focused on the neural response elicited by the cueing syllable and subsequent neutral impulse during WM retention, as both could reactivate ordinal rank information^39^. All trials were labeled according to the continuous ordinal rank (1-9) of the cueing syllable, regardless of the identity of the syllables. Trials in which the cueing syllable was associated with identical local and global ranks (i.e., the 1^st^, 5^th^, 9^th^ syllable in the whole sequence) were excluded from the analysis, since these trials could not distinguish local and global ranks. The multivariate neural dissimilarity, i.e., Mahalanobis distance between 64-channel EEG response or 204-sensor MEG gradiometer response in each 50-ms time window (5 values for each sensor), between neural responses to syllables of different ranks were then calculated, yielding the neural representational dissimilarity matrix (neural RDM; Figure 1D, upper left, purple matrix).

To fit the neural RDM, a number of model RDMs (Euclidean distance) were generated based on the representational geometry model by sampling all possible pairing of LGA and LGS (Figure 1D, upper right, black matrix). Using a regression analysis (linear mixed model), we evaluated which model RDM, i.e., which parameter of the representational geometry model, best fitted the neural RDM, with fixed effects indicating across-subject effects and random effects indicating individual effects. This regression analysis was conducted at each time point, resulting in time-resolved model fitting performance, quantified by negative log-likelihood, as a function of LGA and LGS (Figure 1D, lower panel). The model parameter leading to the lowest negative log-likelihood was selected as the best model parameter (red asterisk) quantifying the neural representational geometry of syllable sequences in WM.

### 2-D Hierarchical organization of syllable sequences in WM

Thirty-two human subjects participated in Experiment 1 with 64-channel EEG activities recorded. Subjects reported the global (96.59% ± 0.37%) and local rank (97.16% ± 0.40%) with high accuracy. Figure 2A plots the time-resolved model fitting results (fixed effect in linear mixed model) throughout the retention period. Around 300 ms after the cueing syllable, the neural response was significantly explained by the parametric representational geometry model (300 – 420 ms, p_max_ = 0.022, FDR corrected). Critically, the best model within the time range (dotted box) supports the 2-D Hierarchy hypothesis (LGA = 109°, 95% CI: 78°∼161°; LGS = 0.9, 95% CI: 0.57∼1.48; fixed effect beta = 0.003, t(1150) = 6.374, p < 0.001).

**Figure 2.**
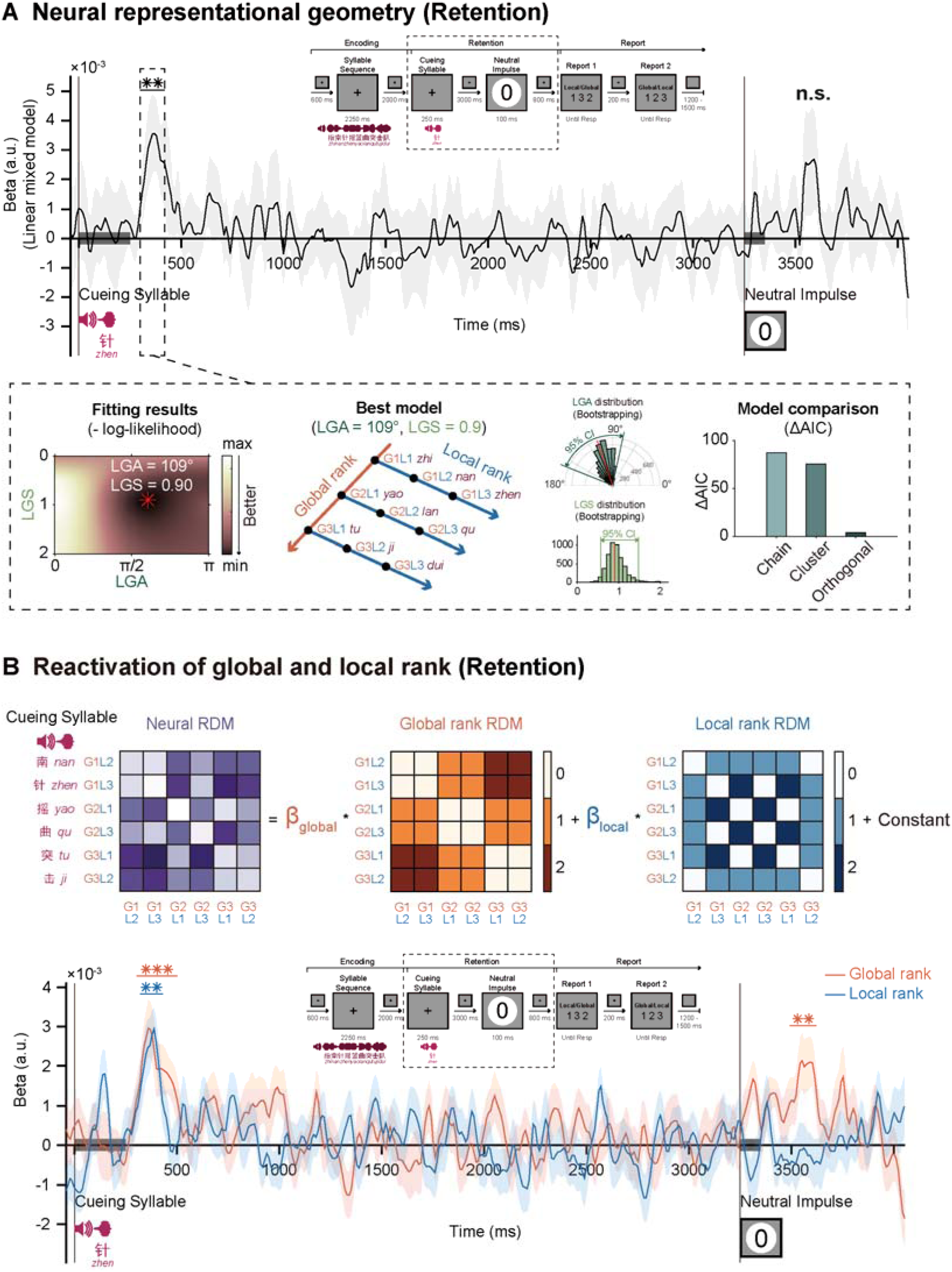
Experiment 1 (N = 32; EEG recordings). (A) Upper: time-resolved performance of the best fitting model (fixed effect in linear mixed model) during WM retention. **: time points with significant p-values (p < 0.05) over three consecutive points after FDR correction. The gray shaded area illustrates the 95% CI for the fixed effect, as determined by the linear mixed model. Lower: best model within the significant time range (dotted box) after the cueing syllable (from left to right): model fitting performance (negative log-likelihood) as a function of LGA and LGS (best model highlighted with red asterisk, LGA = 109°, LGS = 0.90); neural representational geometry of the best model; LGA distribution (N = 5000, Bootstrapping with LGS fixed at 0.90) and LGS distribution (N = 5000, Bootstrapping with LGA fixed at 109°); comparison between the best model and the three hypotheses with fixed parameters (1-D Chain: LGA = 0°, LGS = 1/3;1-D Cluster: LGA = 0°, LGS = 1/4; 2-D Orthogonal: LGA = 90°, LGS = 1). (See Figure S1 and Figure S2). (B) Neural decoding of global and local rank information during WM retention. Upper: neural RDMs were regressed using global rank RDM (orange) and local rank RDM (blue). Lower: grand average (N = 32, mean ± SEM) decoding performance of global (orange) and local (blue) rank information as a function of time (nonparametric sign-permutation test, cluster-based permutation correction, p < 0.05; ***: p < 0.001; **: p < 0.05).

To further confirm these findings, Akaike Information Criterion (AIC) values for the best model (LGA = 109°, LGS = 0.9) were compared against those of the three hypotheses: 1-D Chain (LGA = 0°, LGS = 1/3), 1-D Cluster (LGA = 0°, LGS = 1/4 as a representative value), and 2-D Orthogonal (LGA = 90°, LGS = 1 as a representative value). Consistently, the Hierarchy hypothesis demonstrated superior performance, exhibiting the smallest ΔAIC compared to the AIC value of the best model (Chain: ΔAIC = 103.641; Cluster: ΔAIC = 90.516; Orthogonal: ΔAIC = 5.741). Crucially, the model comparison results hold at the individual level as well (Figure S1A; Orthogonal vs. Chain: t(31) = 3.696, p < 0.001, Cohen’s d = 0.653, 95% CI: 0. 000452 ∼ 0.00156; Orthogonal vs. Cluster: t(31) = 3.359, p = 0.002, Cohen’s d = 0.594, 95% CI: 0.000322∼0.00132; paired t-test). In contrast, the neural response after the neutral impulse was not significantly explained by the parametric representational geometry model (Figure 2A, 290 – 350 ms after impulse onset, p_max_ = 0.154, FDR corrected; see Figure S2A for details).

In addition, we sought to directly decode global and local rank information from neural responses. Specifically, each participant’s neural RDM was regressed with two model RDMs, one based on the global rank and the other based on the local rank (Figure 2B, upper). The cueing syllable indeed triggered neural encoding of both global and local ranks (permutation test; global rank: 300 – 500 ms, p < 0.001, corrected; local rank: 320 – 430 ms, p = 0.008, corrected), while the neutral impulse only triggered global (240 – 370 ms, p = 0.003, corrected) but not local rank (p_min_ = 0.382, corrected) (Figure 2B, lower).

In summary, these analyses demonstrated that a 1-D syllable sequence was retained in the WM according to the 2-D Hierarchy hypothesis, with separate neural dimensions for the local and global ranks.

### 2-D hierarchical organization of syllable sequences with varied word length

Experiment 1 exclusively presented trisyllabic words. In Experiment 2 (Figure 3A), we varied the word length to introduce more variability and flexibility and test the generality of the 2-D Hierarchy hypothesis. Specifically, the syllable sequence contained a random combination of words with 2 to 4 syllables (Figure 3B). Moreover, instead of presenting syllables at a fixed rate, 0-40 ms temporal jitter was added. Thirty-two subjects participated in Experiment 2 and reported the global (96.77% ± 0.37%) and local rank (97.54% ± 0.28%) with high accuracy.

**Figure 3.**
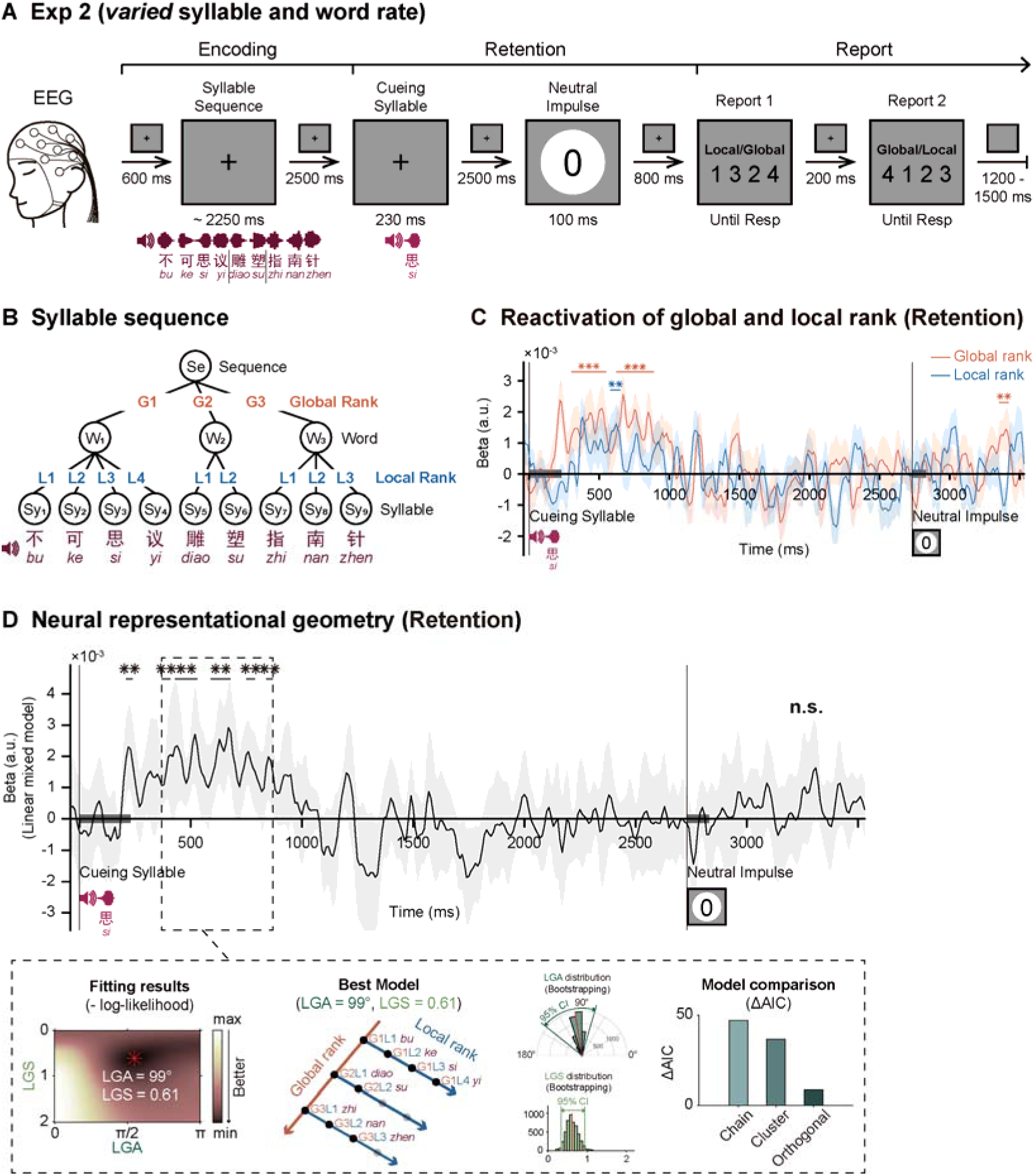
Experiment 2 (N = 32; EEG recordings). (A) Same as Experiment 1 but using syllable sequences with varying word lengths and adding random inter-syllable temporal jitters (0 - 40 ms). (B) Each syllable sequence contained random combination of words encompassing 2 to 4 syllables. (C) Same as Figure 2B lower panel. (D) Same as Figure 2A. (See Figure S1 and Figure S2).

Experiment 2 successfully replicated the results of Experiment 1 (Figure 3D). The neural response elicited by the cueing syllable was significantly explained by the representational geometry model (370 ms – 870 ms; best model fixed effect beta = 0.002, t(1150) = 7.239, p < 0.001), and the fitted model parameter was consistent with the 2-D Hierarchy hypothesis (LGA = 99°, 95% CI: 74° ∼ 141°; LGS = 0.61, 95% CI: 0.34 ∼ 0.94). Model comparison (Chain: ΔAIC = 47.021; Cluster: ΔAIC = 36.412; Orthogonal: ΔAIC = 8.359) and individual-level analysis (Figure S1B; Orthogonal vs Chain: t(31) = 2.161, p = 0.039, Cohen’s d = 0.382, 95% CI: 0.000018 ∼ 0.000623; Orthogonal vs Cluster: t(31) = 1.701, p = 0.099, Cohen’s d = 0.301, 95% CI: −0.000045 ∼ 0.000501; paired t-test) also confirmed the 2-D hierarchical hypothesis. Also, consistent with Experiment 1, neural activity following the neutral impulse was not significantly explained by the representational geometry model (p_max_ = 0.594, FDR corrected; see Figure S2A for details). Finally, as shown in Figure 3C, the cuing syllable elicited both global and local rank information (permutation test; global rank: 300 – 550 ms, 620 – 890 ms, p < 0.001, corrected; local rank: 580 – 650 ms, p = 0.043, corrected), while the neutral impulse only reactivated global rank information (global rank: 620 – 680 ms, p = 0.043, corrected; local rank: pmin = 0.080, corrected). In summary, even when the syllables were grouped into words of different lengths, the 1-D syllable sequence was retained in the WM according to the 2-D Hierarchy hypothesis.

### 2-D Hierarchical organization of syllable sequences is task-irrelevant

In Experiments 1 and 2, subjects were asked to separately report the global and local ranks of the cueing syllable, which encourages the separation of global and local rank information in WM and may drive the 2-D neural representation. To investigate whether the 2-D representation is created automatically or on task demand, we performed Experiment 3, wherein subjects did not need to explicitly calculate the global and local ranks and only reported the continuous rank of the cueing syllable in the whole sequence (1-9) with their MEG activities recorded (Figure 4A). We also removed the neutral impulse since it did not reactivate local rank information in Experiments 1 and 2. Thirty subjects participated in Experiment 3 and reported the continuous rank (96.04% ± 0.63%) with high accuracy.

**Figure 4.**
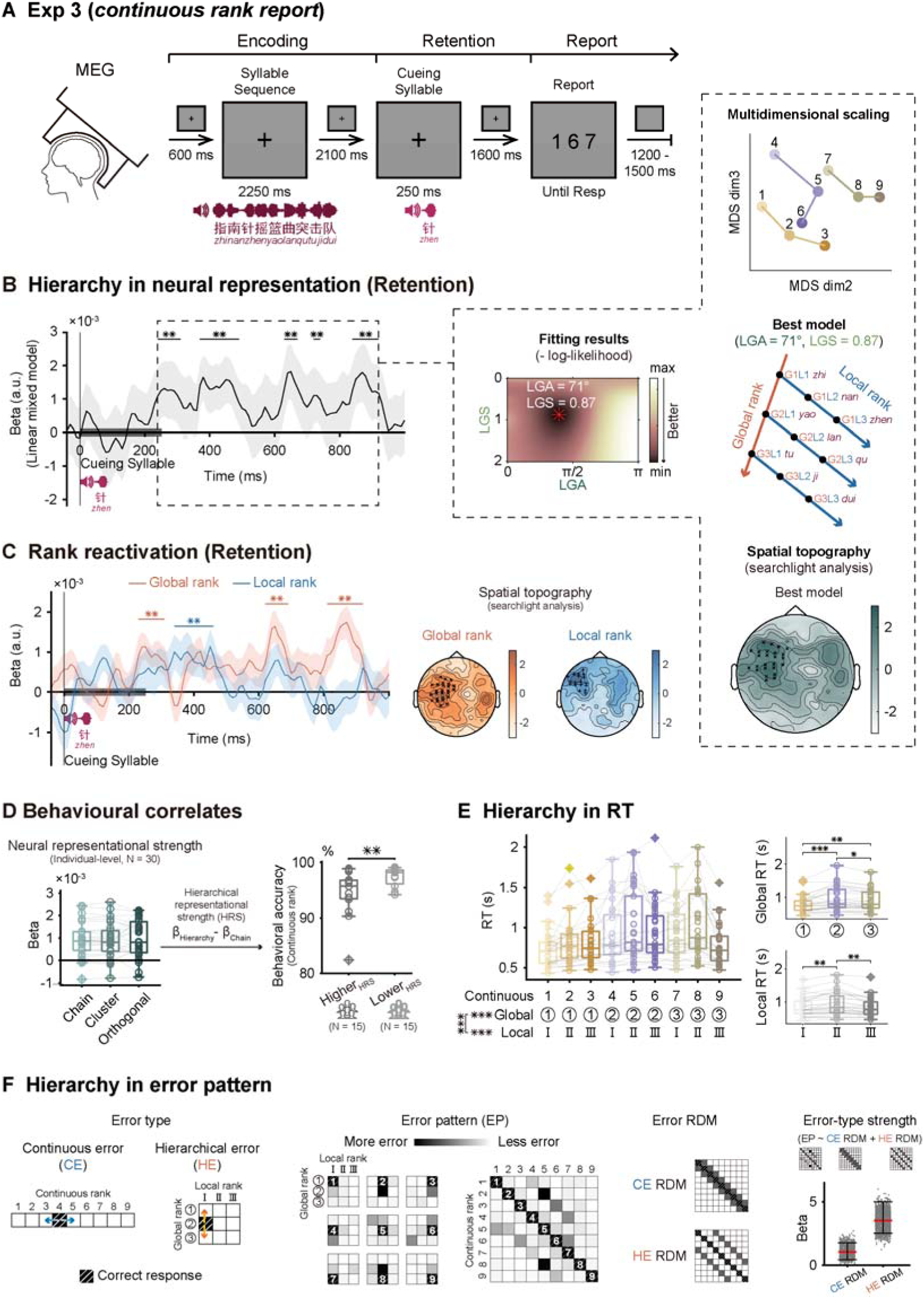
Experiment 3 (N = 30; MEG recordings). (A) Procedure is the same as Experiment 1 but the task is to report the continuous rank (1-9) of the cueing syllable. (B) Same as Figure 2A, adding sensor-level searchlight results for the best neural geometry model (lower panel in the dotted inset; nonparametric sign-permutation test, cluster-based permutation correction, p < 0.05; **: p < 0.05). (C) Left: Same as Figure 2B. Right: sensor-level searchlight results for global (red) and local (blue) rank representations (nonparametric sign-permutation test, cluster-based permutation correction, p < 0.01; **: p < 0.05). (D) Behavioral correlates of neural representational geometry. Left: individual-level regression coefficient of neural RDM to the three hypotheses (1-D Chain, 1-D Cluster, 2-D Orthogonal). Each dot denotes one subject. Subjects were next divided into two groups based on their hierarchical representational strength (HRS) index (*β_orthogonal_ - β_chain_*). Right: continuous rank report accuracy of Higher_HRS_ and Lower_HRS_ groups (independent t-test, **: p < 0.05). Each dot denotes one subject. (E) Hierarchical structure in RT patterns. Left: RT as a function of cueing syllable’s continuous rank (two-way repeated ANOVA, ***: p< 0.001). Right-upper: RT as a function of global rank by marginalizing local rank; Right-lower: RT as a function of local rank by marginalizing global rank (paired t-test, ***: p < 0.001; **: p < 0.05; *: p < 0.1). Each dot denotes one subject. (F) Hierarchical structure in error pattern. Left: Illustration of two types of error based on two hypotheses. Continuous error (CE): reported error based on 1-D chain hypothesis (blue arrow). Hierarchical error (HE): reported error based on 2-D hierarchy hypothesis (orange arrow). Middle-left: observed error patterns pooled across subjects. The 9 small 3×3 matrices denote error distribution for each of the 9 continuous ranks (1-9) organized in 2-D form. The 9×9 matrix is the combination of the 9 3×3 matrices, with each line corresponding to each 3×3 matrix. Middle-right: Error RDMs for CE (upper) and HE (lower). Right: Regression of observed error patterns with RDM_CE_ and RDM_HE_ (Bootstrapping, N = 5000, 95% confidence interval). Each dot represents one bootstrapping iteration.

Experiment 3 replicated the main findings of Experiments 1 and 2, regardless of the critical task change (Figure 4B). The cueing syllable elicited neural responses that could be well captured by the parametric representational geometry model but in a more sustained manner (240 – 920 ms, best model fixed effect beta = 0.001, t(2428) = 6.919, p < 0.001). The fitted parameter was consistent with the 2-D Hierarchy hypothesis (LGA = 71°, 95% CI: 74° ∼ 141°; LGS = 0.87, 95% CI: 0.41 ∼ 1.39), and the result was confirmed by model comparison (Chain: ΔAIC = 25.729; Cluster: ΔAIC = 25.488; Orthogonal: ΔAIC = 8.107). We further performed a multidimensional scaling (MDS) analysis on the neural RDM, which exhibits a clear hierarchical organization of the syllable sequence in WM (Figure 4B, right dotted box, upper panel). Lastly, we conducted a searchlight analysis (permutation test; corrected) in the sensor space and revealed one cluster within left frontal-parietal channels that are involved in the 2-D geometry representation of hierarchical structures (Figure 4B, right dotted box, bottom panel). This suggests that instead of arising from dissociated regions encoding global and local ranks separately, the 2-D geometry originates from neural operations within higher-level brain regions (also see Figure 4C, right panel).

Consistent with Experiments 1 and 2, both global and local ranks were triggered by the cueing syllable (Figure 4C, left panel; permutation test; global rank: 230 – 310 ms, p = 0.014, 620 – 690 ms, p = 0.023, 810 – 920 ms, p = 0.003, corrected; local rank: 340 – 460 ms, p = 0.003, corrected). The searchlight analysis further confirmed that global and local ranks are processed in overlapping frontal-parietal areas (Figure 4C, right panel; permutation test; corrected).

Overall, even in a continuous-report task that encourages the subject to store the 1-D sequence of syllables as it is, the brain still automatically reorganizes the 1-D syllable sequence into a 2-D neural representational geometry in WM.

### 2-D Hierarchical organization in WM behavior

On top of MEG evidence, behavioral responses during the continuous-rank report task also supported the Hierarchy hypothesis. First, the reaction time (RT) calculated for each of the 9 cueing positions showed that primacy and recency effects were observed both at the local and the global scales, i.e., shorter reaction time for the first and last word and shorter reaction time for the first and last syllable within each word (Figure 4E). Two-way repeated ANOVA analyses supported the contribution of both global and local ranks to RTs (global rank: F(1.364,39.569) = 15.050, p < 0.001, partial *η*^2^ = 0.343; local rank: F(1.358,39.390) = 13.098, p < 0.001, partial *η*^2^ = 0.311; interactions: F(3.083,89.405) = 15.758, p < 0.001, partial *η*^2^ = 0.352).

The reported error profiles also support the 2-D Hierarchy hypothesis. As illustrated in Figure 4F (left), the 1-D Chain and 2-D Hierarchy hypotheses predict different error patterns, which were separately referred to as the Continuous error (CE) and the Hierarchical error (HE). Specifically, if the sequence is retained as a 1-D chain as the task required, a syllable tends to be confused with its neighboring syllables in the sequence, e.g., the 4^th^ syllable would be reported as the 3^rd^ or 5^th^ syllable (Figure 4F, left, CE, blue arrow). In contrast, if syllables are organized based on a 2-D hierarchical representation, a syllable can also be confused with its neighbors along the global rank axis, i.e., the 4^th^ syllable confused with the 1^st^ or 7^th^ syllable (Figure 4F, left, HE, orange arrow). The observed errors displayed a pattern consistent with HE profile (Figure 4F, middle-left, Error pattern). To further quantify the results, we built two RDMs based on CE and HE hypotheses (Figure 4F, middle-right, Error RDM), and regressed the overall error pattern by CE and HE RDMs (Figure 4F, right). The results support the greater contribution of Hierarchical structure to behavior (Bootstrapping test; *β_CE_* = 1.025, 95% CI: 0.440 ∼ 1.769; *β_HE_* = 3.523, 95% CI: 2.507 ∼ 4.998).

### Neural representational geometry predicts WM behavior

Finally, we examined whether the neural representational geometry was related to WM behavior in individual subjects. Since the continuous-report task in Experiment 3 motivated subjects to maintain a 1-D chain neural representation, a 2-D hierarchical neural geometry, although observed at the population level, could in principle adversely affect the continuous-report behavioral performance. We thereby hypothesized that individuals showing 1-D chain neural geometry commensurate with task requirements demonstrated higher accuracy on the continuous-report task than those showing 2-D hierarchical neural geometry.

To test this hypothesis, we regressed individual neural RDM with the model RDMs derived from the 1-D Chain and 2-D Orthogonal hypotheses (Figure 4D, left panel), based on which we computed a hierarchical representational strength (HRS) index for each participant, defined as the difference between the regression beta values of the 2-D Orthogonal and 1-D Chain hypotheses. Consistent with our hypothesis, subjects with higher HRS demonstrated lower report accuracy (94.64% ± 4.22%) than subjects with lower HRS values (97.45% ± 1.73%) on the continuous rank report task (Figure 4D, right panel; independent t-test; t(28) = - 2.389, p = 0.024, Cohen’s d = 0.872, 95% CI: - 5.22% ∼ - 0.402%).

Taken together, subjects automatically reorganize syllable sequences into a 2-D hierarchical neural representational geometry in WM even when the reorganization is unfavorable to the task. The findings also further substantiate the strong link between neural representational geometry in WM and memory behavior.

## Discussion

It has long been proposed that WM organize information into hierarchically embedded chunks, yet the underlying neural implementation remains unknown. In three human EEG/MEG experiments, we provide converging evidence that hierarchically organized structures underlying a 1-D sensory sequence are represented by 2-D neural representational geometry in WM, whereby each item in the sequence is reorganized along two dimensions, separtely corresponding to chunk order and the order of an item within a chunk. The 2-D geometry, arising from parieto-frontal regions, is automatically constructed regardless of task demands and could predict behavior.

Hierarchical organization is found in a wide range of human behaviors, including language, music, knowledge, motor control, memory, decision making and planning^12,13,45–47^. Humans indeed excel at tracking the hierarchical structure embedded in stimuli^48,49^, inferring hierarchically generative rules^50^, and making hierarchically structured plans^14,19^. This hierarchy-based organizational principle offers a compelling solution to balance capacity limits and flexible operations in WM, such as compressing information storage, decreasing interference between levels, and boosting retrieval efficiency by top-down signals^18,19,51,52^. Here, even in a task that encourages a 1-D representation by reporting continuous 1-D rank, behavioral measurements (i.e., RT and reported-error profile) exhibit hierarchical structures, consistent with previous findings^21–23,25,53–55^, and global/local ranks could be separately decoded from neural activities. These findings add substantial evidence for the intrinsic hierarchical nature of WM.

Most importantly, we demonstrate that this hierarchical reorganization in WM is neurally implemented through 2-D geometry rather than 1-D clustered geometry, although both operations could achieve WM chunking. Previous modeling works propose that items in a hierarchical sequence are associated with a global and a local index^26–28^, advocating a 2-D representation in mental space. Our findings support this view and provide novel insight into its neural implementations. Notably, hierarchically organized sequences do not necessarily correspond to 2-D geometry in neural space. For example, brain activities track information at multiple hierarchical levels^48,56,57^, yet this could not reveal how those levels are related to each other. Here, the 2-D neural geometry constitutes an abstract, unified neural space that can reveal the internal chunk organization of items, regardless of the content, and this neural geometry can support the representation of chunks of varying lengths (i.e., Experiment 2). The close association with WM behavior further confirms its central role in structure-based WM operations. In fact, the 2-D factorized encoding strategy is proposed to minimize inference between representational dimensions^40,42,58^. Finally, the 2-D neural geometry arises from concentrated parietal-frontal regions, largely excluding the possibility that the two representational dimensions are mediated by dissociated brain areas, e.g., lower- and higher-level regions^6,9,59^. Instead, our findings are more consistent with recent monkey recordings revealing that PFC neural populations contain nearly orthogonized neural subspaces for ranks in a non-hierarchical simple sequence^37^. Together, we postulate that this 2-D neural geometry provides a scaffold for complex sequences containing multiple hierarchical levels via the development of new axes, which together form an embedded structure.

Different from works examining the encoding period when items are presented^43,44,48,49^, here we focused on WM retention, a stage that presumably better reflects internal WM organizational rules. To this end, we presented triggering events (i.e., cueing syllables and neutral impulses) during retention to elicit order-specific neural responses^60–64^. Notably, unlike high-precision intracranial recordings, time-resolved WM information could not be robustly extracted using noninvasive electrophysiological recordings ^65,66^. Accordingly, memories are posited to be retained in activity-silent states and only reactivated by transient perturbations^62–64^. The current study does not aim to address the active vs. silent nature of WM, but leverages the efficacy of cuing syllables in triggering the phase-locked, strong neural response that carries rank information^39,67–69^. Moreover, previous studies demonstrated sustained rank-related activities^34,70^, while reactivations here tend to be more transient. This discrepancy might be due to whole-sequence production tasks employed previously^34,70^. Furthermore, neural replay facilitates sequence memory, structure representation, and novel inference, or in a broad sense, the formation of cognitive maps^71–74^. We hypothesize that replay, as a 1-D sequence pattern, might arise from temporal sampling of an underlying 2-D neural geometry, considering their noted dynamic differences: fast and compressed for replay and relatively slow and stable for neural geometry.

We leveraged linguistic materials to examine neural correlates of hierarchical structure in WM. Could the 2-D neural geometry just reflect the syllable associations formed in long-term semantic memory? Syllables within words (local ordinal rank) might be partially impacted by their semantic relationship, as explicitly modeled in our 1-D cluster hypothesis. Meanwhile, the global ordinal rank could not arise from the semantic relationship at all, since words are arbitrarily selected and concatenated in each trial. Moreover, phonetic similarity between syllables also falls short of explaining the findings, as syllables possess distinct pronunciations and could not contribute to their neural similarities. Finally, as 2-D neural geometry could be generalized to sequences with unpredictable chunk boundaries, i.e., combinations of varying word lengths, temporal prediction also fails to account for the findings.

Taken together, we demonstrate 2-D geometrical representations of complex sequences in WM and their impact on behavior. The findings illustrate how the brain automaticlly reorganizes incoming information into an appropriate structure that facilitates information storage and retrieval, and would provide potential insights into a wide range of domains given WM’s involvement in almost any cognitive function.

## Methods

### Subjects

Thirty-two (18 females and 14 males, mean age 20.8, range 18-27 years), thirty-two (17 females and 15 males, mean age 21.3, range 18-27 years), and thirty (11 females and 19 males, mean age 21.2, range 18-27 years) native listeners of Mandarin Chinese with no history of neurological or psychiatric disorders were recruited in Experiment 1, Experiment 2, and Experiment 3, respectively. No subject participated in more than one experiment. All subjects had normal or corrected-to-normal vision and none of them had any known auditory disorders. All subjects gave written informed consent prior to taking part in the experiments. The experiments received approval from the Departmental Ethical Committee of Peking University. Subjects received compensation for their participation in the form of either 60 RMB per hour or course credits.

### Apparatus of EEG experiments (Experiment 1 and Experiment 2)

The whole experiment was performed via Psychophysics Toolbox-3^75^ for MATLAB (MathWorks) using custom scripts. Visual stimuli were presented on a 32-inch Display++ LCD screen, operating at a resolution of 1920 x 1080 pixels and a refresh rate of 120 Hz. Auditory stimuli (i.e., syllable sequences and cueing syllables) were generated in advance using custom MATLAB codes. During the experiment, these auditory stimuli were delivered to the subjects via a Sennheiser CX300S earphone that was connected to an RME Babyface pro external sound card. Subjects’ head positions were secured using a chin rest, which was positioned 75 cm away from the screen. Subjects’ responses for the tasks were collected using a standard QWERTY keyboard.

### Apparatus of MEG experiment (Experiment 3)

The whole experiment was also controlled via Psychophysics Toolbox-3^75^ for MATLAB (MathWorks) using custom scripts. Visual stimuli were projected onto a 32-inch rear projection screen, positioned 75 cm away from the subjects, using a projector. The projector operated at a resolution of 1920 x 1080 pixels and a refresh rate of 60 Hz. The pre-generated auditory stimuli (i.e., syllable sequences and cueing syllables) were delivered to the subjects via an air-tube earphone that was connected to an RME Babyface pro external sound card. Subjects’ responses were collected using a response pad.

### Auditory stimuli

In the current research, auditory stimuli were syllable sequences and cueing syllables. All stimuli were presented in Mandarin Chinese, where each character corresponds to a single syllable and combinations of characters (i.e., syllables) form words conveying specific meanings. In both Experiment 1 and Experiment 3, syllable sequences were formed by concatenating three trisyllabic words. In Experiment 2, syllable sequences were constructed by randomly selecting and concatenating three words from a pool comprising disyllabic words, trisyllabic words, and quadrisyllabic idioms. We ensured that each syllable sequence did not contain any syllables with the same pronunciation. Although the syllable sequences could be structured into three multisyllabic words, each of the constituent syllables was independently synthesized using the Neospeech synthesizer (http://www.neospeech.com/, the male voice, Liang). The synthesized syllables had a duration range of 168 to 397 ms (mean 250 ms) in Experiment 1, 168 to 400 ms (mean 251 ms) in Experiment 2, and 155 to 401 ms (mean 250 ms) in Experiment 3. To achieve uniformity, all syllables in Experiment 1 and Experiment 3 were adjusted to a consistent length of 250 ms, either by truncating the syllable or adding silence at the end. Those syllables in Experiment 2 were adjusted to 230 ms. The final 25 ms of each syllable were smoothed by a cosine window. Upon the synthesis of constituent syllables, they were successively concatenated without any temporal gaps, forming the syllable sequences for both Experiment 1 and Experiment 3. For the syllable sequences in Experiment 2, a random temporal jitter ranging from 0 to 40 ms was introduced between two successive syllables. At present, all syllable sequences have been generated. Given the acoustic independence of constituted syllables, the hierarchical structure of syllable sequences (syllable-word-sequence) could only be extracted through semantic knowledge (Figure 1A), not prosodic cues.

In this setup, each syllable within the syllable sequence carries two pieces of structural information: local rank information (the ordinal rank of the syllable within its multisyllabic word) and global rank information (the ordinal rank of the multisyllabic word within the entire sequence). All texts and sound files of the syllable sequence and cueing syllables are available here (https://osf.io/drzuy/?view_only=4649d27224464d2ea98d3ba501d8443c) **Experimental procedure (Experiment 1)**

The initiation of a trial was signaled by the presentation of a black cross (0.9° in visual angle), positioned centrally against a gray background (RGB = (128,128,128)). This cross remained in place throughout the entirety of a trial, with the exception of during the neutral impulse presentation and the report screen (Figure 1C). Subjects were instructed to maintain their gaze on the central cross and minimize eye blinking throughout each trial. During the encoding phase of the working memory task, subjects were presented with pre-generated syllable sequences via auditory delivery. The task required subjects to memorize these sequences in the order they were presented. During the maintenance phase, two types of trigger events, an auditory cueing syllable, which was one of the syllables delivered during the encoding phase, and a visual neutral impulse (visual angle is 18°), were employed to probe the neural representation of hierarchical structure. The rationale is that is that each syllable is associated with specific hierarchical structure information during the encoding phase, based on the current syllable sequence. As a result, when a syllable is presented during the maintenance period, its associated hierarchical structure information could be reactivated. The visual neutral impulse, meanwhile, is designed to perturb the working memory network, thereby reactivating the maintained information. These two trigger events were proved efficient in probing sequential temporal structures (1^st^, 2^nd^, and 3^rd^) in our previous research^39^. The subjects’ task was to report both the global and local rank of the cueing syllable. The order in which these ranks were reported was randomized for each trial and was displayed on the report screen. For each report, subjects were instructed to use their index, middle, or ring fingers of their right hand to press the “j”, “k”, or “l” keys, respectively. Notably, to prevent motor response preparation during the maintenance phase, the correspondence between the ranks (1^st^, 2^nd^, and 3^rd^) and the response keys (“j”, “k”, or “l”) was randomly assigned for each trial, as indicated on the report screen. In Experiment 1, we delivered only those syllables as cueing syllables that corresponded to inconsistent global and local ranks (i.e., global 1 and local 2 (G1L2), G1L3, G2L1, G2L3, G3L1, and G3L2). This was done to avoid scenarios where the global and local ranks of a cueing syllable are identical. In such cases, subjects could complete the experimental task by representing a single rank information, without specifying whether it pertains to a global or local rank. This could potentially introduce ambiguity in discerning whether the neural representation is related to global or local hierarchical information. Experiment 1 consisted of a total of 432 trials. To mitigate fatigue, a mandatory break of at least one minute was instituted after every 30 trials.

### Experimental procedure (Experiment 2)

Experiment 2 closely mirrors the design of Experiment 1, with the key distinction being that the syllable sequences in Experiment 2 exhibit irregular hierarchical structures (Figure 3B; details see Auditory stimuli). Furthermore, certain timing parameters were modified to manage the overall duration of the experiment (Figure 3A). The response keys were also expanded to four keys, with the “j”, “k”, “l”, and “;” keys corresponding to the index, middle, ring, and little fingers of the right hand, respectively. Experiment 2 comprised a total of 486 trials.

### Experimental procedure (Experiment 3)

The hierarchical structure of the syllable sequences in Experiment 3 is identical to that of Experiment 1, both of which are constructed by concatenating three trisyllabic words. However, the specific syllable sequences used in Experiment 3 were newly generated. Most importantly, the behavioral task was to report the continuous rank of the cueing syllable within the syllable sequence, which ranged from 1-9, at the end of each trial (Figure 4A). Contrary to tasks that involve reporting the global and local ranks of the cueing syllable in Experiment 1 and 2, the task in Experiment 3 does not encourage any hierarchical structuring of the syllable sequence explicitly. Therefore, the Experiment 3 could eliminate the possibility that findings from Experiment 1 and 2 are solely confined to the hierarchical rank report task. In addition, Experiment 3 employed syllables corresponding to all continuous ranks (1-9) as the cueing syllable. Meanwhile, we chose to retain only cueing syllable as the trigger event in Experiment 3 given its proven efficiency in reactivating hierarchical structures in both Experiment 1 and Experiment 2. Experiment 3 comprised a total of 459 trials.

### EEG acquisition and pre-processing (Experiment 1)

The EEG signals were acquired using a 64-channel EasyCap (Brain Products, Herrsching, Germany). The data were recorded using two BrainAmp amplifiers (Brain Products, Herrsching, Germany) and the BrainVision Recorder software (Brain Products, Herrsching, Germany) at a frequency of 500 Hz. Throughout the entire EEG recording process, the impedance of all electrodes was maintained below 10 kΩ. The recorded EEG data were subsequently pre-processed offline utilizing FieldTrip^76^ in MATLAB 2022a. The data were segmented into epochs extending from 200 ms prior to the onset of the syllable sequence to 500 ms following the onset of the report screen. These epochs were then baseline-corrected using the mean activity from 50 ms to 100 ms prior to the onset of the syllable sequence as the baseline for subtraction. Following this, the data were re-referenced to the average value across all channels, down-sampled to a frequency of 100 Hz, and subjected to a bandpass filter within the 1 Hz to 30 Hz range. Independent component analysis employing FastICA algorithm was conducted to eliminate components associated with eye movement and other artifacts, such as bad channels and heartbeat. The remained components were back-projected into the EEG channel space for subsequent analysis.

### EEG acquisition and pre-processing (Experiment 2)

The acquisition of EEG data in Experiment 2 was carried out in the same manner as in Experiment 1. The pre-processing procedure was also largely identical, with the exception that epochs were extracted from 2500 ms prior to the onset of the cueing syllable to 700 ms following the offset of the neutral impulse in Experiment 2.

### MEG acquisition and pre-processing (Experiment 3)

Neuromagnetic data were acquired using a whole-head MEG system with 204 planar gradiometers and 102 magnetometers (Elekta Neuromag system, Helsinki, Finland) in a magnetically shielded room. The MEG experiment was divided into ten blocks, each followed by a brief break. Before the commencement of each block, the position of the subject’s head was estimated using index coils placed at four points on the head. This position was then compared to the initial position recorded at the start of the experiment to ensure that any head movement did not exceed 4mm throughout the experiment. Magnetic field strength was sampled at a frequency of 1000 Hz. The recorded MEG data were subsequently pre-processed offline utilizing MNE-Python tools^77^. Initially, the data underwent de-noising and motion correction using the Maxfilter Signal Space Separation method. Subsequently, the data were band-pass filtered to a range of 1-40 Hz and downsampled to a frequency of 100 Hz. Epochs were then extracted, specifically from 600 ms prior to the onset of the syllable sequence to 600 ms following the onset of the report screen. Finally, ICA employing fastICA algorithm was conducted to eliminate components associated with eye movement and heartbeat.

### Representational similarity analysis (RSA) based on parametric representational geometry model

#### (1) Neural representational dissimilarity matrix (RDM)

Each entry in the neural RDM represents the Mahalanobis distance between neural representations of two hierarchical ranks, which were reactivated by cueing syllable. (Figure 1D, upper left). In the computation of neural RDM, each trial was labeled based on the hierarchical rank associated with the cueing syllable. For instance, consider a three trisyllabic word sequence to be memorized as “ZhiNanZhenYaoLanQuTuJiDui”, where the cueing syllable is “Zhen”. Given that the “Zhen” is the third syllable of the first word, this trial would be labeled as global 1 local 3, abbreviated as G1L3. Furthermore, a subsampling of trials was performed to guarantee that within the sampled trials, no cueing syllables shared the same pronunciation. This subsampling procedure was implemented to ensure that the computed neural dissimilarity across different hierarchical ranks was not influenced by the recurrence of the same cueing syllable pronunciation across various ranks. Meanwhile, during the trial subsampling process, we ensured an approximately equal number of trials for each hierarchical rank to preserve comparability across conditions. For subjects with the fewest trials, each hierarchical rank included 28 trials. Finally, the calculation of the neural RDM incorporated both spatial and temporal features^78^. Specifically, at each time point, signals from all sensors (64 EEG channels for Experiments 1 and 2, and 204 gradiometers for Experiment 3) within a forward 50 ms time window (5 values, one for each of five 10 ms windows) were included as spatiotemporal features. This resulted in 320 features (i.e., 64*5) for the EEG experiments and 1020 features (i.e., 204*5) for the MEG experiment at each time point.

We utilized an 8-fold cross-validated approach to compute the neural RDM^38,79^. The previously subsampled trials were partitioned into eight folds, ensuring an approximately equal distribution of trials for each hierarchical rank within each fold. Subsequently, one fold was taken as testing dataset, while the remaining seven folds constituted training dataset. The condition-specific spatiotemporal neural pattern was computed by averaging the trials that corresponded to identical hierarchical ranks within the training dataset. Subsequently, the Mahalanobis distance was computed between the spatiotemporal neural activity of each individual trial in the testing dataset and each condition-specific spatiotemporal neural pattern that was derived from the training dataset. Finally, the original Mahalanobis distances was adjusted by subtracting the mean distance between each testing trial and all training conditions for each testing trial (i.e., mean-centered). The above procedure was iteratively performed until each of the eight folds had served as testing dataset. Upon completion, we obtained a neural representational dissimilarity matrix of dimensions corresponding to the number of trials by the number of conditions (i.e., hierarchical ranks). Finally, distances of trials corresponding to identical hierarchical ranks were averaged, resulting in a condition-by-condition neural RDM. The entire procedure, from trial subsampling to the generation of the final condition-by-condition neural RDM, was performed 300 times. The resulting 300 neural RDMs were then averaged to yield the final neural RDM at each time point.

#### (2) Parametric representational geometry model

It is proposed that the representational geometry for abstract hierarchical structure can be projected onto a two-dimensional plane (Figure 1D, upper right). This plane is defined by two axes, each representing a distinct level of rank information: one corresponds to high-level global ranks, and the other to low-level local ranks. The specific geometry can be quantified using two parameters: local-global angle (LGA) and local-global scale (LGS). The LGA represents the angle between representational axes of local and global ranks, while the LGS denotes the ratio of unit lengths of these two axes. Upon determining a pair of LGA and LGS values, a specific geometry can be defined. Subsequently, the model distance (d; Euclidean distance in the two-dimensional plane) between each pair of hierarchical ranks can be calculated based on this defined geometry, resulting in a model representational dissimilarity matrix (model RDM). Through the systematic sampling of parameters LGA (ranging from 0° to 180° in 1° increments) and LGS (ranging from 0 to 2 in 0.01 increments), we were able to construct a series of hypothetical neural geometries. Each of these geometries is associated with a corresponding model RDM.

#### (3) Estimation of the genuine representational geometry

Upon gathering all candidate model RDMs, each model RDM were regressed with the neural RDM via a linear mixed model to ascertain which model RDM that best fits the neural RDM, as indicated by the smallest negative Log-Likelihood value for the model fitting. The selected model RDM then permitted the deduction of the neural representational geometry of hierarchical structure in working memory (Figure 1D, lower). Lastly, it is crucial to highlight that systematic sampling of LGA and LGS guarantees the conventional Chain form (LGA = 0°, LGS = 0.25), Cluster form ((LGA = 0°, LGS = 0.33), and Hierarchy form (Orthogonal form; LGA = 90°, LGS = 1) are all encompassed within all model RDM candidates.

The above RSA based on the parametric representational geometry model was performed at each time point during WM maintenance period. This allowed us to identify the best model at each time point, along with its fixed effect strength (β) and level of significance (Figure 2A time course; Figure 3D time course; Figure 4B time course). Given the substantial number of time points, we applied False Discovery Rate (FDR) correction to adjust for multiple comparisons. For visualization only, the fixed effect strength (β) time courses were smoothed with a Gaussian-weighted window (window length = 40 ms).

In order to summarize neural geometry for hierarchical structure representation in working memory, the neural RDMs across those time intervals (Figure 2A upper dotted box; Figure 3D upper dotted box; Figure 4B left dotted box) that demonstrated a significant fixed effect in the linear mixed model fitting were averaged. The averaged neural RDM were then utilized for subsequent analyses (Figure 2A lower dotted box; Figure 3D lower dotted box; Figure 4B right dotted box): 1) The previously introduced RSA based on the parametric representational geometry model was used to illuminate the parametric model fitting results and determine the geometry for the averaged neural RDM within significant time intervals; 2) The Akaike Information Criterion (AIC) values for the obtained best geometry were compared against those of the three conventional hypotheses: 1-D Chain, 1-D Cluster, and 2-D Orthogonal, to evaluate which hypothesis best fit the neural data. This allowed us to determine which of these predefined geometric structures provided the most accurate representation of the underlying neural organization. and 3) In Experiment 3, multi-dimensional scaling (MDS) was employed to directly visualize the neural representational geometry from the recorded signals. To achieve this, we first recalculated the neural representational dissimilarity matrices (RDMs). This involved calculating the condition-specific spatiotemporal neural activities using all subsampled trials and then computing the pairwise Mahalanobis distance between these activities. This computation resulted in symmetric neural RDMs with zero diagonals, which are suitable inputs for the classical MDS method. Finally, by applying classical MDS, we projected the averaged symmetric RDM across all subjects and significant time points into a three-dimensional space, ultimately deriving the final neural geometry.

### Model-based neural decoding analyses

To directly investigate the reactivation patterns of both global and local rank information for cueing syllables during maintenance period, we implemented model-based neural decoding analyses (Figure 2B; Figure 3C; Figure 4C left). Specifically, for each subject individually, the time-resolved neural RDM was regressed using two predictors: the global rank RDM and the local rank RDM. Each regression coefficient (β) time courses were smoothed with a Gaussian-weighted window (window length = 40 ms). A regression coefficient (β) significantly exceeding zero signifies that corresponding information is represented in the neural activities. We employed a non-parametric sign-permutation test^80^ to conduct statistical analyses on the regression coefficients. Specifically, the sign of regression coefficients for each subject at each time point was subjected to 100,000 random flips, which process allowed the generation of null distribution of the population mean β. The p-value of the observed population mean β was then estimated from this null distribution. In order to adjust for multiple comparisons over time, a cluster-based permutation test was conducted, with a cluster-forming threshold of p < 0.05.

### Searchlight analyses

A searchlight analyses was performed to identify MEG sensors (gradiometers) involved in representing the 2-D Hierarchical geometry, global rank, and local rank. For each gradiometer, we computed a neural multivariate RDM incorporating neural activities from this gradiometer and its neighbors (averaging 8.33 neighbors, range: 4-11) within a 50-ms time window as features at each time point. Next, the neural RDMs across significant time windows identified for 2-D Hierarchy (240-920 ms; Figure 4B, left) were averaged and regressed with the best model RDM (LGA = 71° and LGS = 0.87). This was performed for each gradiometer, yielding the spatial distribution of neural representation strength. Similar procedures were applied to both global and local ranks, within the corresponding significant time windows (global: 230-920 ms, local: 340-460 ms; Figure 4C, left). Model-based neural decoding analyses (Figure 2B, upper) were then conducted on the averaged neural RDMs to derive the spatial topography for each rank. Finally, cluster-based permutation tests (Monte Carlo method for cluster-based permutation; cluster-forming threshold p < 0.05) were conducted to identify gradiometers with significantly higher regression coefficients compared to the group median value.

### Correlations between brain and behavior

To quantify the predictive power of neural representational strengths on behavioral responses in the continuous rank report task, we conducted a median split analysis in Experiment 3. Specifically, for each subject, we first averaged the neural representational dissimilarity matrix (RDM) within the identified significant time window (Figure 2B left, dotted box). This averaged RDM was then individually regressed against the three predefined RDMs corresponding to the Chain model, Cluster model, and Orthogonal model. Subsequently, we computed the hierarchical representational strength (HRS) by calculating the difference in regression coefficients between the Orthogonal (task irrelevant) and Chain (task relevant) models. Finally, subjects were divided into two groups based on a median split of their HRS values. To assess the relationship between HRS and behavior, we compared the behavioral response accuracy of the higher HRS group with the lower HRS group using a paired t-test.

### Model-based behavioral decoding analyses

In order to examine whether the hierarchical organization of the syllable sequence is reflected in the patterns of error responses, we carried out model-based behavioral decoding analyses (Figure 4F). Erroneous responses from all subjects were collectively aggregated. For each continuous rank (1-9), we calculated the distribution of erroneous responses, which we denote as the error pattern (EP). Regression analysis was applied to the EP using two predictors. The first, continuous error RDM, represents a scenario where a syllable’s rank could be mistaken as its adjacent ranks in a one-dimensional chain (e.g., the fourth syllable could be mistaken as the third or fifth). The second predictor, hierarchical error RDM, encapsulates instances where a syllable’s rank could be confused with its adjacent ranks along the global rank dimension in a two-dimensional hierarchical form (e.g., the fourth syllable could be mistaken as the first or seventh). A significant regression coefficient (β) implies the existence of a corresponding organization within the error pattern. We employed a bootstrapping method (N = 5000) for the statistical analysis of the regression coefficients, ensuring that the current results were not disproportionately influenced by a single subject. Meanwhile, given our focus on incorrect responses, the diagonal elements of the matrices were omitted from the analyses. What’s more, it’s worth noting that mistaking the fourth item for the fifth could also be a condition where diffusion occurs along the local rank dimension in a two-dimensional hierarchical form. However, we currently classify this scenario as a continuous error, which results in an underestimation of hierarchical error strength and an overestimation of continuous error strength.

## Data availability

All the raw data will be made available upon acceptance (https://osf.io/drzuy/?view_only=4649d27224464d2ea98d3ba501d8443c).

## Code availability

The code to replicate our main analyses will be made available upon acceptance (https://osf.io/drzuy/?view_only=4649d27224464d2ea98d3ba501d8443c).

## Acknowledgments

This work was supported by the National Science and Technology Innovation STI2030-Major Project 2021ZD0204100 (2021ZD0204103 to H.L.), National Natural Science Foundation of China (31930052 to H.L.), and China Postdoctoral Science Foundation (2023M740124 to Y.F.). The authors thank the Center for MRI Research at Peking University in Beijing, China, for assistance with data acquisition. We would like to thank Dongning Liu, Qiming Han, Jiaxin Gao for their help during the data collection, and Jian Li for helpful comments on data analysis.

## Author information

### Authors and Affiliations

School of Psychological and Cognitive Sciences, Peking University, Beijing, China Ying Fan, Muzhi Wang, Huan Luo

PKU-IDG/McGovern Institute for Brain Research, Peking University, Beijing, China Ying Fan, Muzhi Wang, Huan Luo

Beijing Key Laboratory of Behavior and Mental Health, Peking University, Beijing, China

Ying Fan, Muzhi Wang, Huan Luo

Key Laboratory for Biomedical Engineering of Ministry of Education, College of Biomedical Engineering and Instrument Sciences, Zhejiang University, Hangzhou, China

Nai Ding

### Contributions

Y.F. and H.L. originally conceived the study. Y.F. performed the experiments. Y.F. and M.W. analyzed the data. Y.F., N.D., and H.L. wrote the paper.

### Corresponding authors

Correspondence to Nai Ding or Huan Luo.

### Ethics declarations

The authors declare no competing interests.

