## Supplemental Figures for "2-D Neural Geometry Underpins Hierarchical Organization of Sequence in Human Working Memory"

Supplementary Figure 1

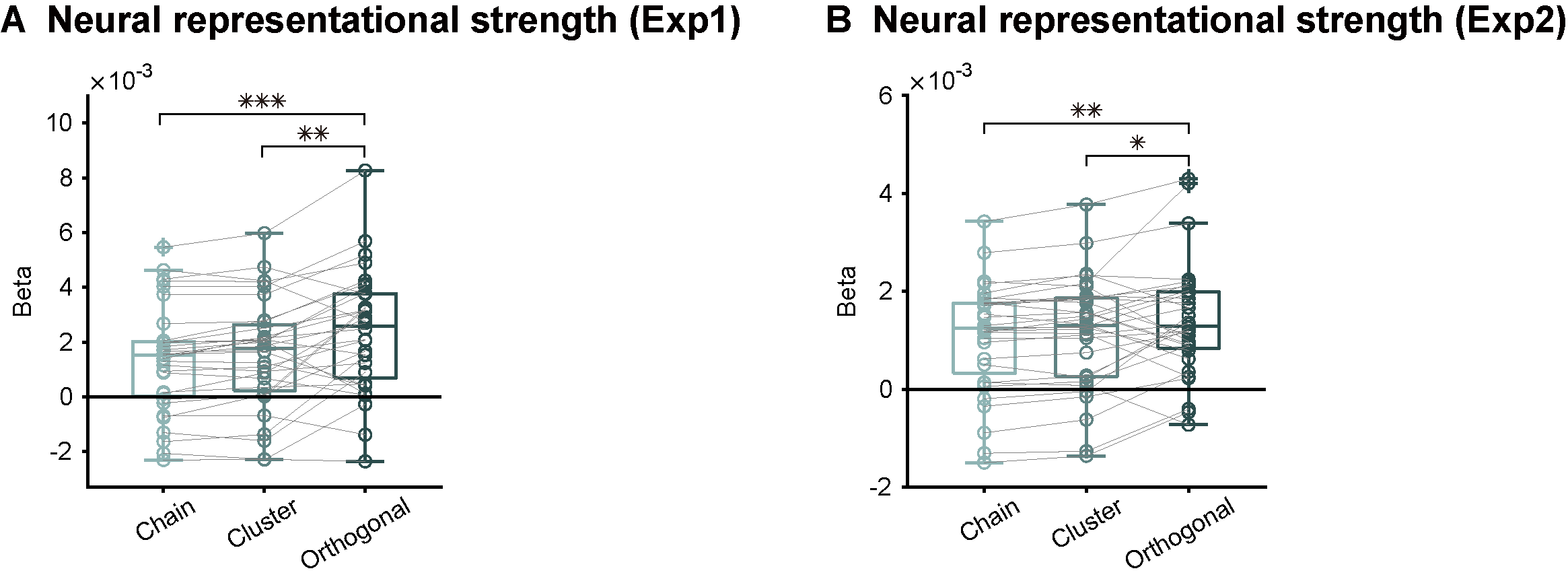

**Figure S1. Neural representational strength in Experiment 1 (related to Figure 2A) and Experiment 2 (related to Figure 3D). (A)** Individual-level regression coefficient of neural RDM within the significant time range (dotted box in Figure 2A) to the three hypotheses (1-D Chain, 1-D Cluster, and 2-D Orthogonal) in Experiment 1. Each dot denotes one subject. Paired t-test, ***: p < 0.001, **: p < 0.05, *: p < 0.1. **(B)** Same as A but for Experiment 2.

Supplementary Figure 2

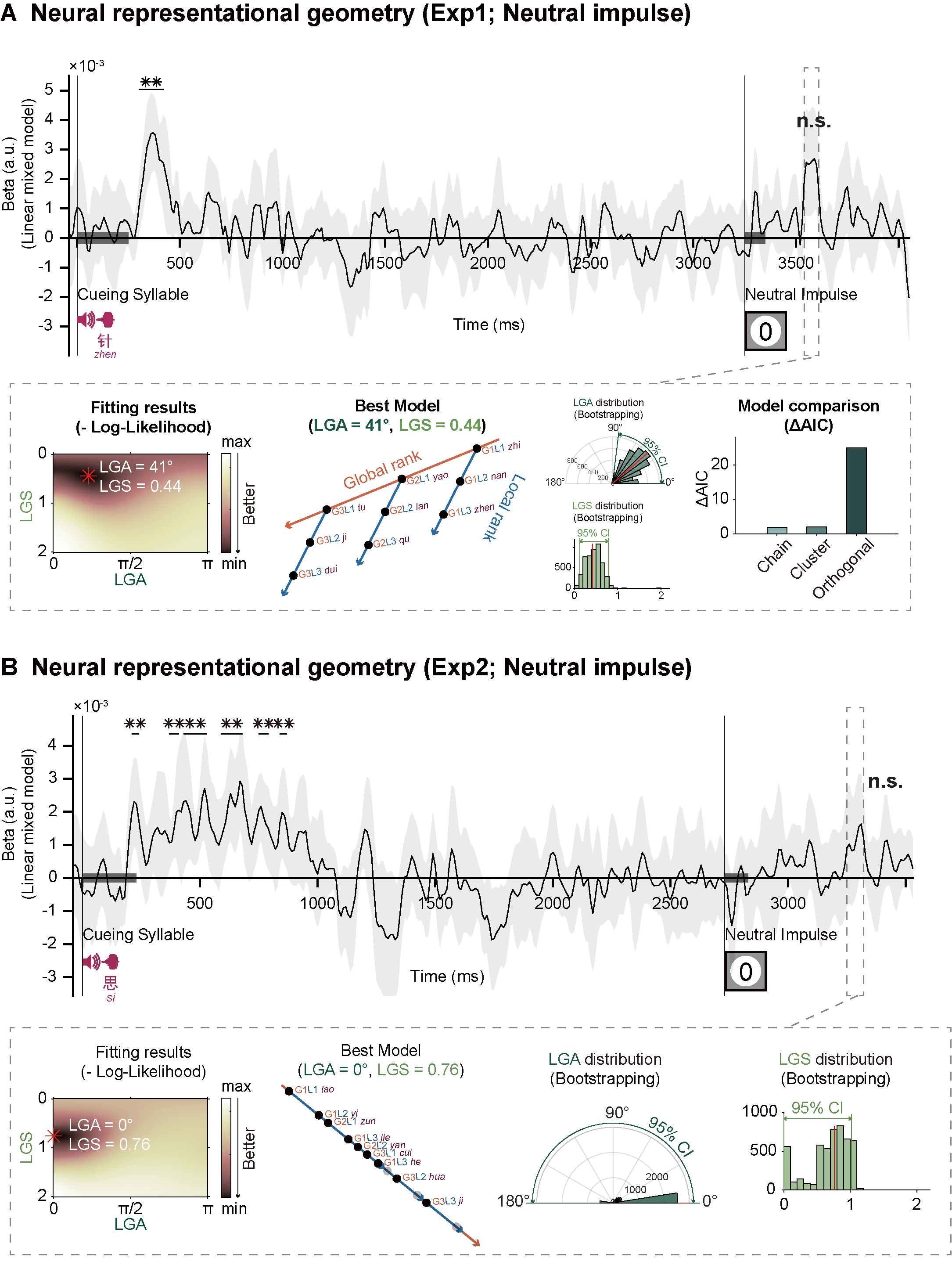

**Figure S2. Neural representational geometry after neutral impulse in Experiment 1 (related to Figure 2A) and Experiment 2 (related to Figure 3D)**. **(A)** Upper: time-resolved model fitting performance (fixed effect in linear mixed model) of neural response during WM retention for Experiment 1. **: time points with significant p-values (p < 0.05) over three consecutive points after FDR correction. Lower: best model within the time range with a high model fitting performance (dotted box; p_max_ = 0.027, uncorrected) after the neutral impulse (from left to right): model fitting performance (negative log-likelihood) as a function of LGA and LGS (best model highlighted with red asterisk, LGA = 41°, LGS = 0.44); neural representational geometry of the best model; LGA distribution (N = 5000, Bootstrapping with LGS fixed at 0.44) and LGS distribution (N = 5000, Bootstrapping with LGA fixed at 41°); comparison between the best model and the three hypotheses with fixed parameters (1-D Chain: LGA = 0°, LGS = 0.33;1-D Cluster: LGA = 0°, LGS = 0.25; 2-D Orthogonal: LGA = 90°, LGS = 1). **(B)** Same as A but for Experiment 2. The maximum and minimum uncorrected p-values associated with the time-resolved model fitting performance within the dotted box are 0.417 and 0.0203, respectively. Given the poor model fitting performance here, comparison between the best model and the three hypotheses was excluded.
